# The Male Default Prevails in Biomedical Research: Sex Inclusion in Nature (2025)

**DOI:** 10.64898/2026.03.03.709344

**Authors:** Ashlyn Swift-Gallant, Liisa A.M. Galea, Lindsay S. Cahill

**Affiliations:** Department of Psychology, Memorial University of Newfoundland, St. John’s NL, Canada; Centre for Addiction and Mental Health, University of Toronto, Toronto, ON Canada; Department of Chemistry, Memorial University of Newfoundland, St. John’s NL, Canada

**Keywords:** Sex as a biological variable (SABV), Sex as a discovery variable, sex inclusion, sex reporting, sex-disaggregated analysis, male-default model, biomedical research, editorial policy, Nature journal

## Abstract

Despite longstanding recognition of sex as a biological variable, its integration into biomedical research remains inconsistent. Numerous publishers have introduced policies to improve reporting and inclusion of sex and gender, including *Nature*, which requires authors to complete a Life Science Reporting Summary documenting sex inclusion. Here, we evaluated the effectiveness of these policies by examining sex inclusion and reporting practices in all original research articles involving humans, vertebrates, or cell lines published in Nature in 2025 (N=513). Nearly two-thirds of articles included both sexes (62.7%); however, inclusion was often nominal. Of these articles reporting inclusion of both sexes, 33% did not maintain inclusion across experiments, used markedly unbalanced sex ratios (≥2:1), or alternated between male- and female-only experiments. Another 45.5% of these articles reporting inclusion of both sexes did not report sample size by sex, so it cannot be ascertained whether sex inclusion was maintained across experiments or balanced by sex. Single-sex studies accounted for approximately one-fifth of articles. While male-only and female-only studies occurred at similar overall rates, male-only studies were more than four times more likely to address conditions affecting both sexes while female-only studies were more likely to address sex-specific conditions (e.g., ovarian cancer). Only 7% of articles explicitly analyzed sex as a discovery variable for at least some analyses. These findings suggest that transparency-focused reporting summaries alone are insufficient to ensure sex inclusion and/or meaningful analytical integration of sex. As a leading biomedical journal, *Nature* plays a central role in shaping research norms; without stronger editorial expectations, reporting requirements risk reinforcing male-default assumptions rather than advancing rigor and generalizability.

## Introduction

The importance of sex as a biological variable in preclinical and clinical research has been acknowledged in the scientific literature for over 30 years (Berkley, 1992; Healt, 1991). Evaluating sex differences has revealed fundamentally distinct biology across organ systems, from brain to heart to kidney, and across endocrine, immune, metabolic, and microbiome pathways, influencing disease prevalence, clinical presentation, and therapeutic response (Becker & Ahmed, 2025; Gogoes, Langmead, Sullivan et al., 2019; Lee, Eid, Hodges et al., 2025). Yet, as recently as 2019, only ∼5% of studies evaluated in neuroscience and psychiatry journals analyzed sex as a discovery variable (i.e., treating sex as an independent variable), and just over half included both sexes (Rechlin, Splinter, Hodges et al., 2022). This means that nearly half of studies still exclude one sex, and mere inclusion rarely leads to direct comparisons, leaving sex-specific differences unexamined. These gaps reveal a pervasive blind spot that limits mechanistic insight, reproducibility, and translational impact.

Given their influence over funding priorities and publication standards, funding agencies and scientific publishers have been described as “primary change agents” essential for advancing sex and gender integration in research (Johnson & Beaudet, 2013). In recent years, these groups have taken more explicit action to promote the integration of sex and gender in research design, analysis, and reporting. The Canadian Institutes of Health Research, a major Canadian funding body for health research, introduced the expectation of sex and gender-based research design and analysis in grant proposals in 2010 (mandatory in 2019). In 2015, the National Institutes of Health in the United States implemented policies requiring applicants to include sex as a biological variable in grant submissions (National Institutes of Health, 2015). Similar efforts have been implemented at national funding agencies worldwide and have been reviewed in (Hankivsky, Springer & Hunting, 2018; White, Tannenbaum, Klinge et al., 2021). The Animals in Research: Reporting *In Vivo* Experiments (ARRIVE) guidelines were published in 2010, with the goal of improving reporting in animal research including description of the sex of the animals used (Kilkenny, Browne, Cuthill et al., 2010). More than 1,000 scientific journals encourage or require the use of the ARRIVE guidelines (Percie du Sert, N., Hurst, V., Ahluwalia, et al., 2020). In 2013, *Nature* took action by introducing a checklist to increase transparency and reproducibility, including requiring sex and gender reporting (Nature Publishing Group, 2013). In 2016, Sex And Gender Equity in Research (SAGER) guidelines were introduced providing specific recommendations to authors, journal editors, peer-reviewers and publishers for ensuring that sex and gender considerations are appropriately reported in the scientific literature (Heidari, Babor, De Castro et al., 2016). And, in 2022, *Nature Portfolio* declared it was “raising the bar”, requiring authors to report how sex and gender were considered in study design, to identify single-sex studies in titles or abstracts, and to provide sex-disaggregated data for studies involving humans, other vertebrates, or cell lines in four of their journals (*Nature Medicine, Nature Communications, Nature Cancer, Nature Metabolism;* Nature Portfolio, 2022).

However, the implementation of checklists and guidelines alone does not guarantee compliance, nor does it necessarily lead to meaningful changes in reporting practices or study design. In the present study, we assessed sex reporting in all *Nature* research articles published in 2025 (note: gender was not evaluated, as gender was only reported in <1% of studies). We focused on *Nature* because it is representative of a top tier, high impact science journal and aims to publish rigorous peer-reviewed research. The objectives of the study are: 1) Evaluate the sex reporting practices in published outputs from 2025. 2) Determine the proportion of articles that included both males and females and if the single sex studies focused on sex-specific conditions. 3) Assess whether articles considered sex as a discovery variable, defined as analyzing sex as an independent variable rather than a covariate or pooling the sexes.

Based on the implementation of a reporting checklist, we hypothesized that the majority of articles will report sex in either the main text or the Reporting Summary. However, given the historical reliance on male subjects as the default model in biomedical research, we hypothesized that single-sex studies, or studies reverting to single-sex experiments after initially including multiple sexes in the article, would remain common. We also expected that a low number of articles would include males and females in the data presentation and compare the sexes in the statistical analysis.

## Results

### Study characteristics and sex reporting practices

We analyzed 513 original research articles published in *Nature* in 2025 that involved human participants, vertebrate animals, and/or cell lines. Sex was reported in 82.1% of articles (n = 421) (Figure 1). While 17.9% of articles (*n* = 92) did not report sex in either the main manuscript or in the Reporting Summary, it is notable that the number is even higher when considering the main manuscript alone: over 30% of studies (*n* = 155) did not disclose sex in the research article itself.

**Figure 1.**
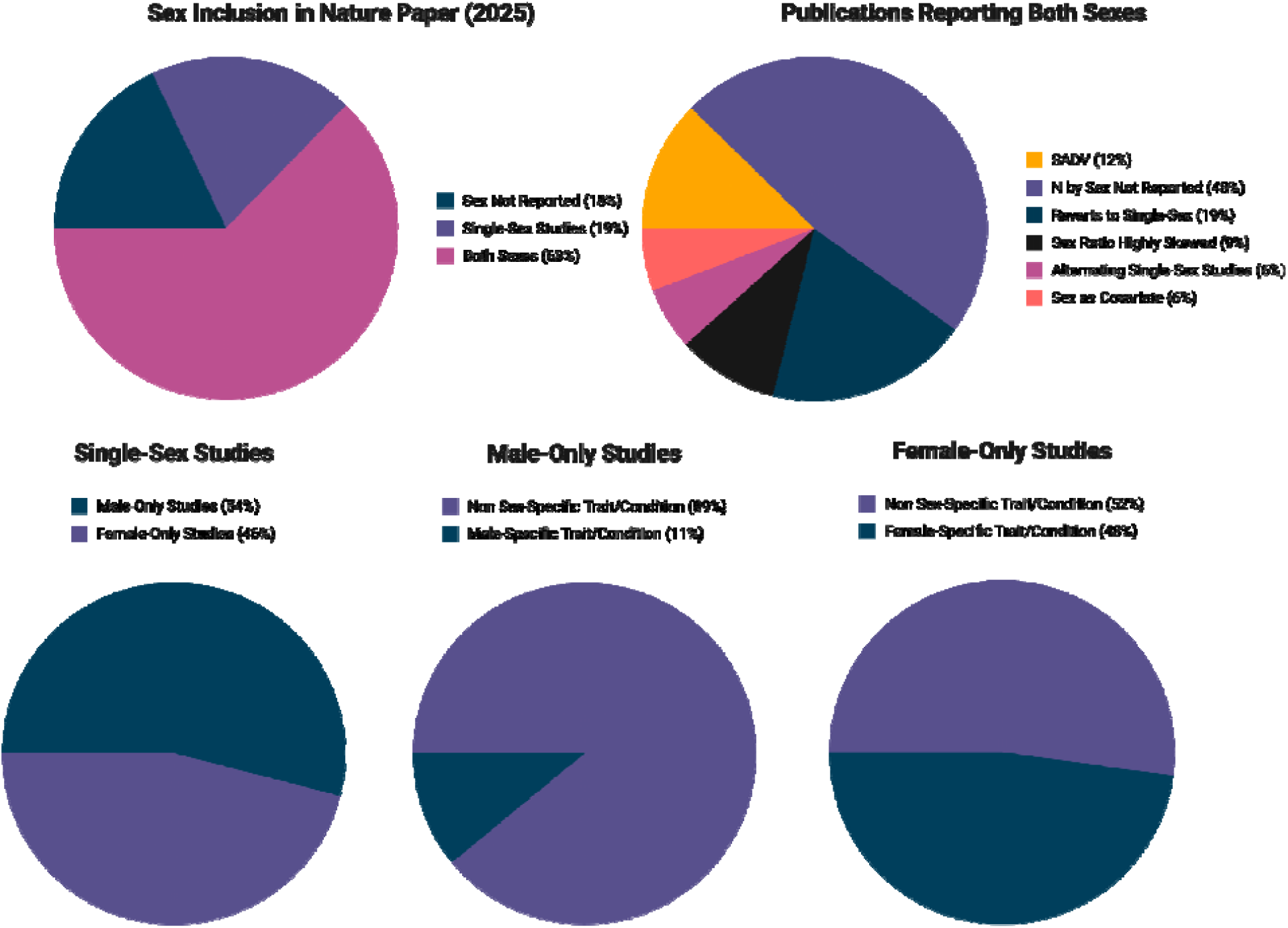
Study characteristics and sex reporting practices. Analysis of 513 original research articles published in Nature in 2025. The first pie chart shows the distribution of sex inclusion: studies including both sexes (62.6%), single-sex studies (19.5%), and studies that did not report sex (17.9%). The second pie chart summarizes sex analysis and reporting practices among studies including both sexes. Only 12% of studies reporting both sexes analyzed sex as a discovery variable for at least some analyses (7% of all studies). Of the remaining studies including both sexes, 48% do not report sample sizes by sex, 19% reverted to single-sex experiments, 9% have highly skewed sex ratios (≥2:1), 6% alternate between male- and female-only experiments, and 6% treat sex as a covariate. The remaining three pie charts depict single-sex studies: female-only (46% of single-sex studies) versus male-only designs (54% of single-sex studies), and, within male-only and female-only studies, the proportions focused on sex-specific versus non–sex-specific conditions.

Most articles (62.6%, *n* = 321) reported inclusion of both sexes. However, among these, 45.5% (*n* = 146) did not provide sex-specific sample sizes in the main manuscript or the Reporting Summary, and/or failed to direct readers to any source where this information could be accessed. An additional 38.3% of articles that reported inclusion of both sexes exhibited one or more of the following patterns: (1) inclusion of both sexes at the outset but reversion to single-sex experiments within the same publication (*n* = 58); (2) alternation between male-only and female-only experiments (*n* = 18); (3) use of skewed samples (*n* = 29; defined as a sex ratio ≥ 2:1); or (4) treatment of sex solely as a covariate (*n* = 18). Thus, over 80% of articles indicating inclusion of both males and females did not do so in a transparent or meaningful manner. Indeed, only 38 articles (11.8% of publications reporting use of both sexes, or 7.4% of all articles) explicitly analyzed sex as a discovery variable in at least *some* analyses. Table 1 summarizes the justifications provided by authors in the reporting summary for not including both sexes and/or for not analyzing data by sex.

**Table 1.**
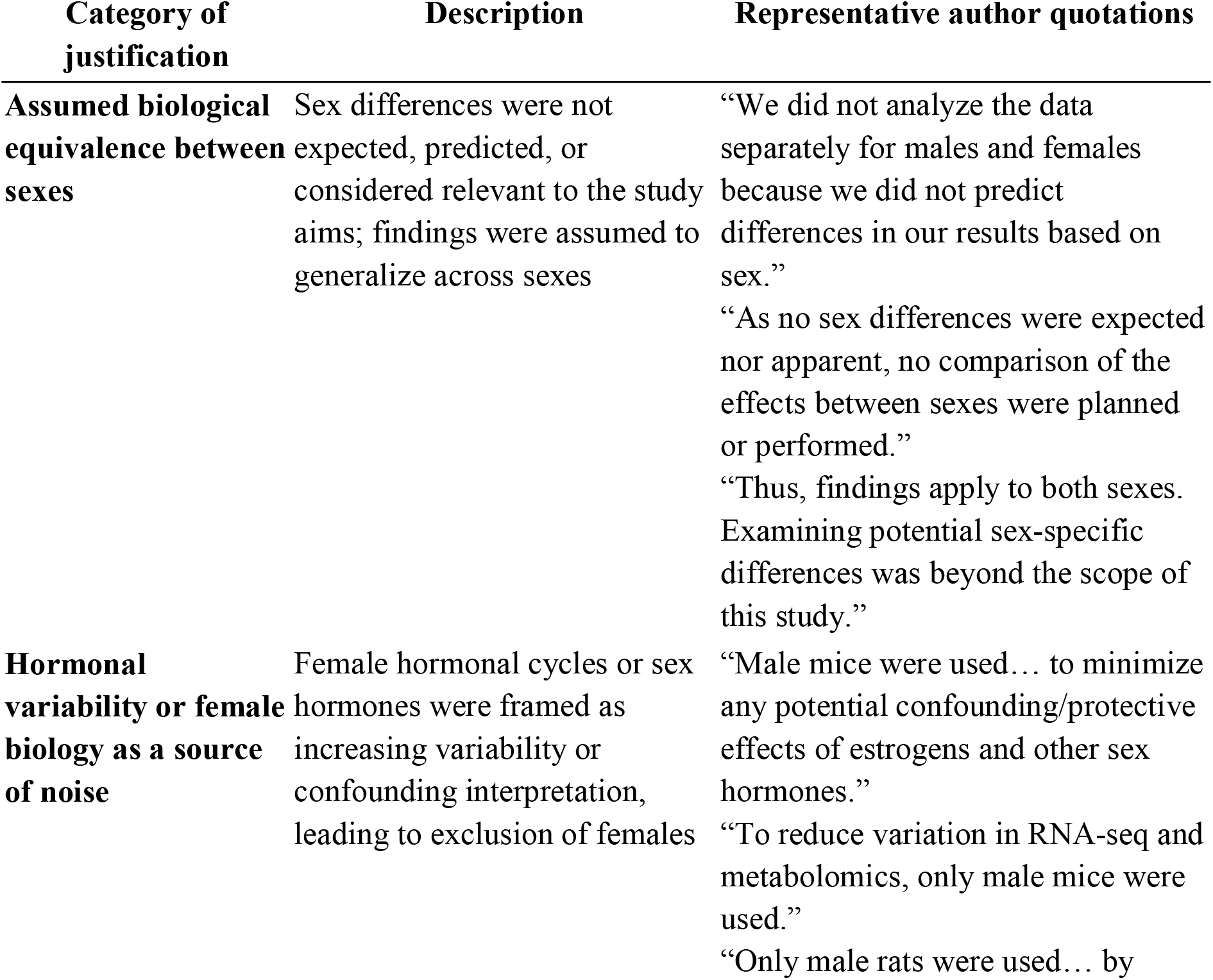
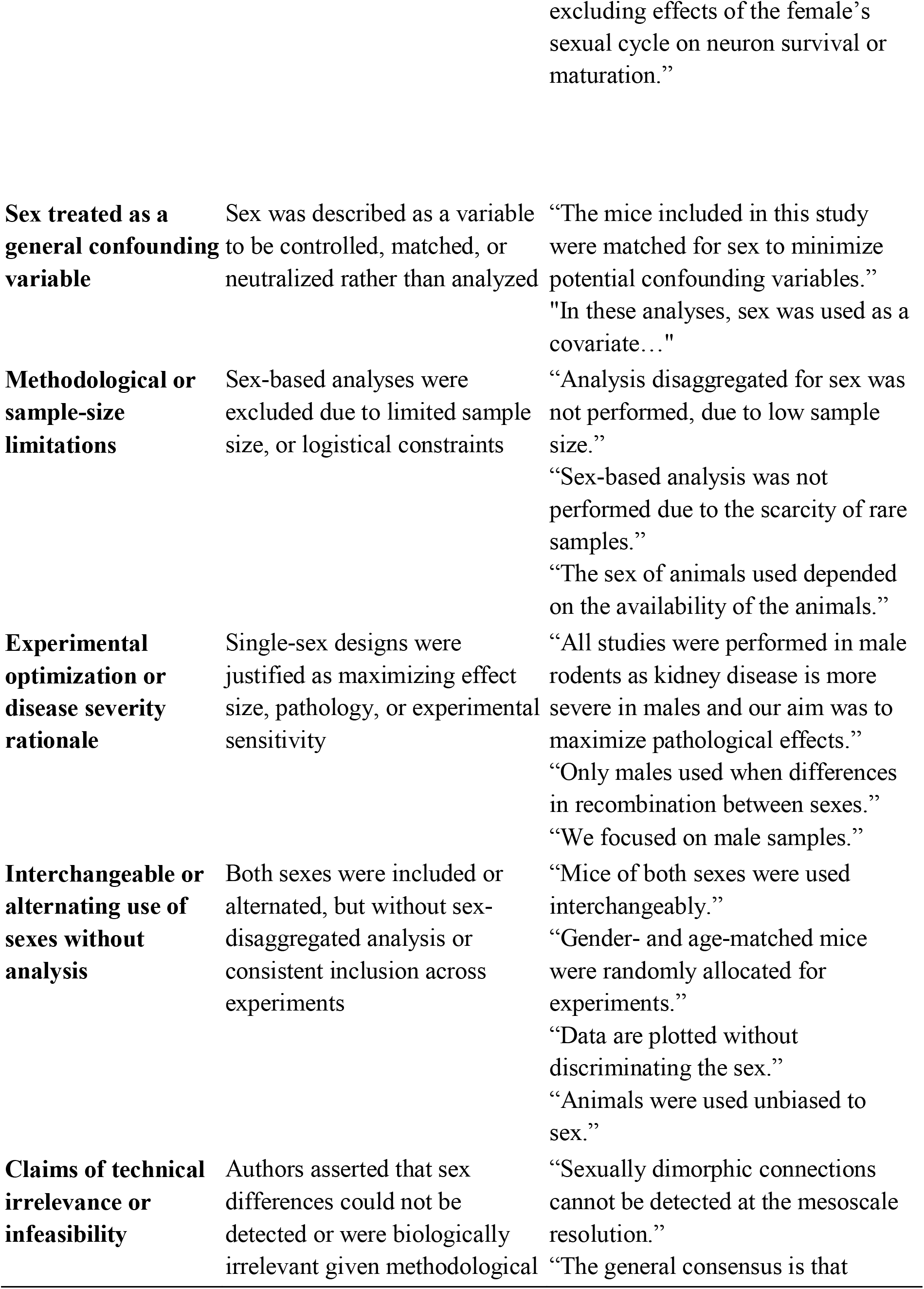
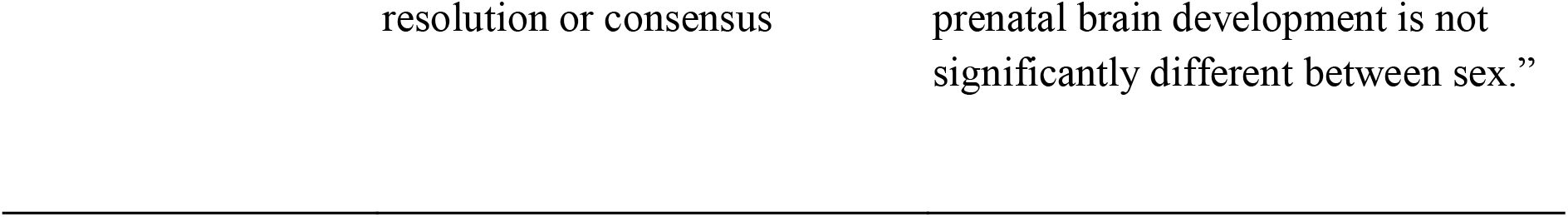
Representative author-reported justifications for single-sex study designs and exclusion of sex-based analyses.

Notably, 19.5% of articles (*n* = 100) employed single-sex designs throughout the publication. Among these, female-only and male-only studies were approximately equally represented: 46 studies (8.97% of all studies; 46% of single-sex studies) used only female subjects or samples, and 54 studies (10.5% of all studies; 54% of single-sex studies) used only male subjects or samples.

Despite similar frequencies of female-only and male-only designs, the scientific focus of these studies differed markedly. Nearly half of female-only studies (47.8%, *n* = 22) investigated female-specific conditions, such as ovarian or endometrial disorders. In contrast, only 11.1% of male-only studies (*n* = 6) focused on male-specific conditions, such as prostate-related diseases. The remaining male-only studies (88.9%) examined conditions that affect both sexes (e.g., aging, neuroscience, liver disease). These findings suggest that female-only designs were largely condition-specific, whereas male-only designs more often excluded females despite studying diseases affecting both sexes. Furthermore, none of the single-sex studies that were not explicitly sex-specific indicated in the title or abstract that they involved only one sex, highlighting a lack of transparency in study reporting.

## Discussion

### Why Nature, and why now

We deliberately focused our analysis on original research articles published in *Nature* in 2025 because *Nature* occupies a uniquely influential position in biomedical research as the flagship journal of the Nature Portfolio. Although *Nature* requires completion of a Life Science Reporting Summary that asks authors to report on sex inclusion, it did not adopt the strengthened, SAGER-aligned reporting and review practices introduced across several other Nature Portfolio journals beginning in 2022. These enhanced measures, including explicit prompts for sex-disaggregated data and pilot efforts to actively engage authors and reviewers, were implemented in journals such as *Nature Medicine, Nature Communications, Nature Cancer*, and *Nature Metabolism*, but not in *Nature* itself (Nature Portfolio, 2022).

It is not clear why these higher standards were not extended to *Nature*, particularly given its central role in setting research norms. One possibility is that existing reporting practices were viewed as sufficient. Our findings, however, suggest otherwise. By examining articles published in *Nature* 2025, we show that transparency-focused reporting summaries alone have not resulted in consistent reporting of sex, inclusion of both sexes, and/or in analysis of sex as a discovery variable. This divergence provides a critical test case for evaluating the limits of reporting requirements in the absence of stronger editorial expectations and enforcement.

### Persistence of the male □ default model and the limits of reporting summaries

Although *Nature* requires authors to report whether both sexes were included in research through its Life Science Reporting Summary, our analysis indicates that male biology continues to function as the implicit default in *Nature* publications. While male□only and female□only studies occurred at similar overall rates, their framing differed sharply: female□only studies were predominantly confined to sex□specific conditions, whereas male□only studies predominated even for traits/conditions affecting both sexes. This pattern reinforces a long□standing conceptual norm in which male biology is treated as broadly generalizable, while female biology is positioned as context□dependent or specialized. Indeed, when resource constraints, limited availability, or methodological considerations are invoked to justify single□sex designs, the choice of sex is not neutral; there is no clear scientific basis for consistently selecting males over females if logistical limitations truly drive design decisions. This persistent avoidance of female□only designs for non-sex□specific conditions implicitly acknowledges the possibility of sex differences - differences that, if examined, would complicate interpretation, require additional investigation, and challenge simplified biological narratives. Prioritizing conceptual simplicity over biological completeness, however, delays the recognition of meaningful sex differences and limits the rigor, generalizability, and translational relevance of research outcomes. A recent example is the case of Lecanemab, an Alzheimer’s disease therapy that received FDA approval in 2023. Lecanemab was originally reported to slow cognitive decline by 27% overall in a Phase□3 trial; however, a secondary analysis conducted years later (Andrews, Ducharme, Chertkow et□al., 2025) revealed a striking sex difference: male participants experienced a statistically significant 43% slowing of decline, while females showed only a 12% improvement. This finding, masked in the original pooled analysis, illustrates how ignoring sex can obscure clinically meaningful differences and potentially affect therapeutic decision-making. This is not an isolated case. Across cardiovascular disease, Alzheimer’s, kidney disease, addiction, immunology, and metabolism, biomedical research has historically relied on male subjects and rarely analyzed or reported results by sex, demonstrating that the masking of sex-specific effects, as seen with Lecanemab, reflects a persistent structural norm rather than a one-time oversight (Becker & Ahmed, 2025).

The experience of nominal inclusion further demonstrates the limits of *Nature’s* reporting summary as a mechanism for changing practice. Although nearly two□thirds of articles reported including both sexes, the majority of these articles did not report sample sizes by sex, did not maintain that inclusion across experiments, used highly skewed sex ratios, or alternated between male- and female-only experiments. Only 7.4% of studies explicitly analyzed sex as a discovery variable. These practices closely mirror the author□reported justifications summarized in Table 1, including claims of assumed biological equivalence, exclusion of females due to perceived hormonal variability, and framing sex as a confounding variable, showing how reporting requirements without analytical expectations allow male-default assumptions to persist. Importantly, this pattern is not unique to *Nature*. Our prior analyses across biomedical journals (2009-2019) have documented similarly low uptake of sex-based discovery approaches, with only ∼5% of studies using sex as a discovery variable (Rechlin, Splinter, Hodges, et al., 2022). Our 2025 findings from Nature are strikingly similar. We focus on *Nature* not to single it out, but because of its leadership role in setting editorial standards; if stronger enforcement and analytical expectations were implemented at highly visible journals, downstream adoption across the field would likely follow. Indeed, prior analyses have documented a significant negative correlation between journal impact factor and SAGER compliance (Michelson, Al-Abedalla, Wagner et al., 2022), raising the paradox that the journals most influential in shaping scientific norms may be among the least consistent in enforcing sex and gender reporting guidelines.

### Implications for editorial policy and research practice

As the flagship journal of the Nature Portfolio, *Nature* plays a disproportionate role in shaping research norms. Our findings suggest that requiring sex reporting through a summary checklist, while valuable for transparency, is insufficient to drive transparency in sex reporting and/or analytical integration of sex as a discovery variable. Extending enhanced sex and gender reporting expectations to *Nature*, including clearer standards for acceptable justifications of single-sex designs, routine reporting of sample size by sex, and encouragement of sex-disaggregated analyses, could substantially improve rigor and generalizability. Without clearer expectations for analytical integration of sex and gender and/or policies to increase topic diversity of female-only studies, *Nature’s* reporting summaries risk becoming administrative checkboxes that entrench male□default assumptions. If the flagship journal of biomedical science cannot model robust integration of sex in study design and analysis, it undercuts incentives for the entire research community to do so, reinforcing the idea that avoiding complexity, even at the expense of rigor and generalizability, is the path of least resistance in high□impact publishing.

## Methods

A search was conducted of all peer-reviewed, original research articles published in Nature from January 1, 2025 to December 31, 2025 (nature.com/nature/research-articles). All studies were screened and included in the analysis if they involved 1) humans, 2) vertebrates, or 3) cell lines (N=513). The following search terms were used in the full research article: “female”, “male”, “women”, “men”, “sex”, “gender”. The Nature Portfolio Reporting Summary was also searched, including “Reporting sex and gender” in the section on human participants, “Reporting on sex” in the section on animals and other research organisms and “Cell line source(s)” in the section on eukaryotic cell lines.

Articles were coded as follows: 1) included sex in the main text or in the Reporting Summary (“yes”, “no”), 2) study characteristics (“both sexes”, “male-only”, “female-only”), 3) presented sex disaggregated data (“yes”, “no”), 4) discussed sex as a biological variable (“yes”, “no”), 5) if only one sex was used, included the sex in the title or abstract (“yes”, “no”), and 6) the species. The reviewers noted when the sample size was not well balanced by sex and when the area of study was a sex-specific condition. They also noted examples of author justifications for the use of only one sex or exclusion of sex-based analyses.

